# *De Novo* Design of Miniprotein Binders Targeting the Leptin-Binding Site of the Leptin Receptor

**DOI:** 10.64898/2026.11.16.766359

**Authors:** Xue Bai, Zhidan Zhao, Yu Jin, Jianyun Guo, Nam-Chul Ha, Ki Hyun Nam, Yongbin Xu

## Abstract

The leptin receptor (LEPR) regulates appetite, energy balance, glucose homeostasis, and neuroendocrine function and is closely associated with obesity, diabetes, and neurological disorders. Activation of LEPR signaling suppresses appetite and promotes energy expenditure, whereas impaired LEPR signaling and leptin resistance are commonly associated with obesity. Accordingly, modulation of LEPR signaling is a potential therapeutic strategy for obesity and related metabolic disorders. In this study, we report the *de novo* computational design of miniprotein binders targeting the leptin-binding site within the cytokine receptor homology 2 (CRH2) domain of LEPR. Computational screening and biochemical characterization identified the miniprotein binder Mini-19484, which adopted a stable α-helical fold and exhibited enhanced resistance to proteolytic degradation. Mini- 19484 bound the LEPR CRH2 domain with high affinity, with a dissociation constant (*K*_d_) of 30 nM. Structural analysis of the predicted model and mutagenesis studies revealed that Arg18 of Mini-19484 serves as a critical interaction hotspot for LEPR binding. The predicted binding mode of Mini-19484 differed substantially from that of natural leptin, providing a distinct molecular scaffold for targeting the LEPR CRH2 domain. These findings provide a valuable framework for developing miniprotein-based therapeutics targeting LEPR.

## 1. INTRODUCTION

Obesity has become a global pandemic [1] and can induce multiple comorbidities, including type 2 diabetes, hypertension, nonalcoholic fatty liver disease, and cardiovascular diseases [2–5]. Obesity is markedly associated with circulating leptin levels [6–8]. Leptin, a key endocrine factor secreted by adipose tissue, regulates feeding behavior, energy expenditure, and multiple metabolic processes [9–12]. Leptin exerts its biological functions through interaction with the leptin receptor (LEPR), which is essential for regulating energy homeostasis, appetite, and body weight [13].

LEPR, a member of the class I cytokine receptor superfamily, comprises an intracellular signaling tail, a single-pass transmembrane domain, and an extracellular ligand-binding domain [14]. The extracellular region possesses a modular architecture characterized by an immunoglobulin-like (Ig) domain, two fibronectin type III (FNIII) domains, and two cytokine receptor homology domains (CRH1 and CRH2) [15] . The CRH2 domain mediates high- affinity interaction with leptin [15]. Leptin binding to LEPR, primarily through the CRH2 domain, promotes receptor activation and downstream signaling that regulate appetite and energy homeostasis [15]. Accordingly, native leptin has attracted considerable interest as a potential therapeutic agent; however, its clinical application remains limited [16]. For example, native leptin has a relatively short circulating half-life and is susceptible to proteolytic degradation, which can limit its therapeutic utility [17, 18]. Moreover, in common obesity, elevated circulating leptin levels often accompany leptin resistance, which is associated with reduced responsiveness to exogenous leptin [19, 20]. Furthermore, metreleptin, an approved recombinant leptin analog, is supplied as a lyophilized formulation and requires refrigerated storage at 2 °C–8 °C, highlighting practical considerations associated with the formulation and storage of leptin-based therapeutics [21–24].

To address these limitations, researchers have also explored alternative strategies targeting LEPR. For example, mibavademab (REGN4461), a fully human agonistic monoclonal antibody, activates LEPR independently of leptin and has been evaluated in a phase 1 clinical trial[25]. Compassionate-use treatment of a patient with atypical partial lipodystrophy and neutralizing antibodies to metreleptin was associated with improvements in triglyceride levels and hepatic steatosis [25]. Researchers have also developed a camelid single-domain antibody to selectively inhibit LEPR–EGFR cross-talk without interfering with canonical LEPR signaling, allowing modulation of leptin-mediated immune responses while preserving its metabolic functions [26]. Despite these advances, antibody-based therapeutics can present challenges related to molecular size, manufacturing complexity, and parenteral administration.

*De novo* computational design is an effective approach for developing high-affinity binding proteins against diverse molecular targets [27, 28]. This strategy has yielded miniprotein binders with picomolar affinity against the SARS-CoV-2 spike protein [29], influenza hemagglutinin [30], and the hACE2 receptor [31], as well as a hyperstable constrained peptide [32], underscoring its broad therapeutic potential. Despite these advances, to our knowledge, *de novo*-designed miniprotein binders targeting the leptin-binding site within the LEPR CRH2 domain have not yet been reported.

In this study, we computationally designed miniproteins targeting the leptin-binding site within the CRH2 domain. We characterized the structural properties and protease resistance of the designed miniproteins and identified an LEPR-binding miniprotein with high binding affinity. Furthermore, structural analysis and mutagenesis identified key residues involved in LEPR binding, and the predicted binding mode of the designed miniprotein was compared with that of natural leptin. These findings show the feasibility of engineering compact and stable LEPR-binding miniproteins and provide a structural and experimental foundation for further development of LEPR-targeting molecules.

## 2. Materials and Methods

### 2.1. Computational design of miniprotein binders

The miniprotein scaffold library was obtained from the Institute for Protein Design repository [33]. Potential binders targeting the LEPR CRH2 domain (residues 430–631) were identified by high-throughput rigid-body docking using PatchDocK [34]. Site specificity was enforced using a secondary filtering script (crmsd) to prioritize miniproteins targeting designated functional hotspots on the LEPR CRH2 domain. The coordinates of the prioritized miniprotein–LEPR complexes were then used as templates for sequence optimization. Miniprotein sequences were redesigned in a fixed-backbone environment of the LEPR receptor using ProteinMPNN [35]. The redesigned sequences were evaluated for their ability to adopt the intended structures using AlphaFold2 [36]. To assess structural fidelity, the predicted models were superimposed onto their corresponding design templates using mTM-align [37, 38]. Only candidates with a root-mean-square deviation (RMSD) of <1.0 Å were retained to ensure preservation of the intended binding orientation. Following this two-step validation, 30 high-quality complex models were selected by visual inspection in PyMOL [39]. The selected LEPR–miniprotein complexes were subsequently subjected to molecular dynamics (MD) simulations using GROMACS [40]. After energy minimization, each complex underwent a 10-ns MD simulation to assess the initial dynamic stability. Predicted binding affinity (*K*_d_) was estimated using PRODIGY [41, 42] before and after the 10-ns MD simulation and was used as an additional criterion for candidate prioritization. Candidates exhibiting favorable dynamic stability and maintained or improved predicted binding affinity were selected for extended 200-ns MD simulations and subsequent experimental validation.

### 2.2. Molecular dynamics simulations

For initial stability screening, 30 redesigned LEPR–miniprotein complexes were subjected to 10-ns MD simulations using GROMACS [40] with the OPLS-AA/L force field [43] at 300 K. Each complex was solvated in a triclinic box containing SPC/E water molecules [44], with a minimum distance of 10 Å between the protein and the box boundary. The systems were neutralized by adding appropriate counterions. Energy minimization was performed using the steepest descent algorithm until the maximum force was <1,0 kJ mol^−1^ nm^−1^. The systems were then equilibrated in two stages: 100-ps NVT equilibration at 37 °C using the V- rescale thermostat, followed by 100-ps NPT equilibration at 1 bar using the Parrinello– Rahman barostat. After equilibration, 10-ns production MD simulations were performed to assess initial complex stability.

Based on the initial MD screening, five LEPR–miniprotein complexes (Mini-9022, Mini- 10457, Mini-12198, Mini-12264, and Mini-19484) were selected for extended all-atom MD simulations. The simulations were performed using GROMACS 2025.4 [40] with an AMBER ff14SB force field [45, 46]. The systems were solvated in a triclinic box containing TIP3P water molecules and simulated at a physiological temperature of 310 K (37 °C). Following appropriate counterion neutralization, energy minimization, and NVT/NPT equilibration steps adapted to these specific parameters, 200-ns production MD runs were executed to thoroughly assess the long-term binding stability of the candidate complexes. Production MD simulations were performed for 200 ns using an integration time step of 2 fs. Covalent bonds involving hydrogen atoms were constrained using the LINCS algorithm. Long-range electrostatic interactions were treated using the particle mesh Ewald method with a cutoff of 1.0 nm. A representative structure of the LEPR–Mini-19484 complex was extracted from the 200-ns production trajectory using RMSD-based clustering with a cutoff of 2.0. Å. The Cα RMSD, radius of gyration, solvent-accessible surface area, and root-mean-square fluctuation (RMSF) were calculated from the MD trajectories using GROMACS analysis tools.

### 2.3. Preparation of the miniproteins

The genes encoding the designed miniproteins were codon-optimized for expression in *Escherichia coli*, synthesized by Sangon Biotech (Shanghai, China), and cloned into the pET-28a expression vector between the NcoI and XhoI restriction sites. The recombinant plasmids were transformed into *E. coli* BL21(DE3) cells. The cells were cultured in 1 L of Luria–Bertani medium containing 50 μg/mL kanamycin at 37 °C until the OD_600_ reached 0.8. Protein expression was induced with 0.5 mM isopropyl β-D-1-thiogalactopyranoside at 30 °C for 8 h. The cells were harvested by centrifugation and resuspended in PBS (137 mM NaCl, 2.7 mM KCl, 10 mM Na_2_HPO_4_, 1.8 mM KH_2_PO_4_, pH 7.4) containing 2 mM β- mercaptoethanol. After sonication, cell debris was removed by centrifugation at 13,000 rpm for 30 min at 4 °C. The resulting supernatant was incubated with 1 mL of Ni-NTA agarose resin (Sangon Biotech, China) with gentle rotation at 4 °C for 1 h and then loaded onto an Gravity-flow column. The resin was washed with PBS (pH 7.4) containing 20 mM imidazole. The target miniproteins were eluted with PBS (pH 7.4) containing 250 mM imidazole. The eluted fractions were pooled and concentrated using a centrifugal concentrator with a 3-kDa molecular weight cutoff (Millipore, USA). During concentration, the buffer was simultaneously exchanged with standard 1× PBS. Finally, protein purity was assessed by SDS-PAGE, and the purified miniproteins were stored at −80 °C until further characterization.

### 2.4. Circular dichroism spectroscopy

The secondary structure and stability of the purified miniproteins were characterized using circular dichroism (CD) spectroscopy. Protein samples were diluted to a final concentration of 0.1–1.0 mg/mL in 20 mM phosphate buffer (12 mM Na_2_HPO_4_ and 8 mM NaH_2_PO_4_, pH 7.4). To assess conformational stability and refolding capability, samples were subjected to three thermal treatments: (i) incubation at 25 °C for 30 min as a room-temperature control; (ii) heating at 100 °C for 5 min followed by immediate cooling in an ice-water bath to 25 °C; and (iii) incubation at 25 °C for an additional 30 min after thermal treatment to assess recovery.

CD spectra were recorded from 190 to 260 nm using a Chirascan CD spectrometer (Applied Photophysics, UK). The background signal was subtracted from all sample spectra for baseline correction.

### 2.5. MicroScale thermophoresis affinity assay

The binding affinity between the LEPR and the designed miniprotein ligands was quantified using microScale thermophoresis (MST). Recombinant LEPR was fluorescently labeled using the Monolith^TM^ Protein Labeling Kit (Cy5). Briefly, 1.4 µL of 14.2 µM LEPR was mixed with Cy5 dye and labeling buffer to achieve final concentrations of 200 nM and 1.2 µM, respectively. After incubation on ice in the dark for 30 min, excess unconjugated dye was removed using a B column (gravity-flow size-exclusion chromatography) pre-equilibrated with assay buffer. The labeled LEPR was eluted with 450 µL of assay buffer after a 550-µL wash and then centrifuged at 12,000 rpm for 10 min at 4 °C to remove dye aggregates. For binding measurements, the miniprotein ligands were serially diluted 2-fold in PBS (pH 7.4) to generate 16 concentrations and mixed in equal volumes (10 µL) with the labeled LEPR to maintain a constant receptor concentration. Samples were loaded into standard treated capillaries and analyzed using a Monolith instrument. Thermophoresis was monitored by measuring changes in normalized fluorescence (*F*_norm_). The dissociation constant (*K*_d_) was determined by nonlinear regression of the experimental data using MO.Affinity Analysis software (NanoTemper Technologies, Germany).

## 3. Results

### 3.1. Revisiting the structures of the CRH2 domain in LEPR

LEPR comprises an extracellular domain (ECD), a transmembrane domain, and an intracellular domain [47]. The ECD represents a potential target for biologic drug development because it enables direct modulation of receptor–ligand interactions using antibodies and peptide-based therapeutics [48, 49]. The LEPR ECD comprises multiple subdomains, including two CRH domains, an Ig domain, and membrane-proximal FNIII domains [15] (Figure 1A). Leptin binds with high affinity to the interface between the two FNIII-like subdomains within the CRH2 domain [50] (Figure 1B). Therefore, miniproteins targeting this interface may competitively inhibit ligand–receptor interactions by blocking receptor binding.

**Figure 1.**
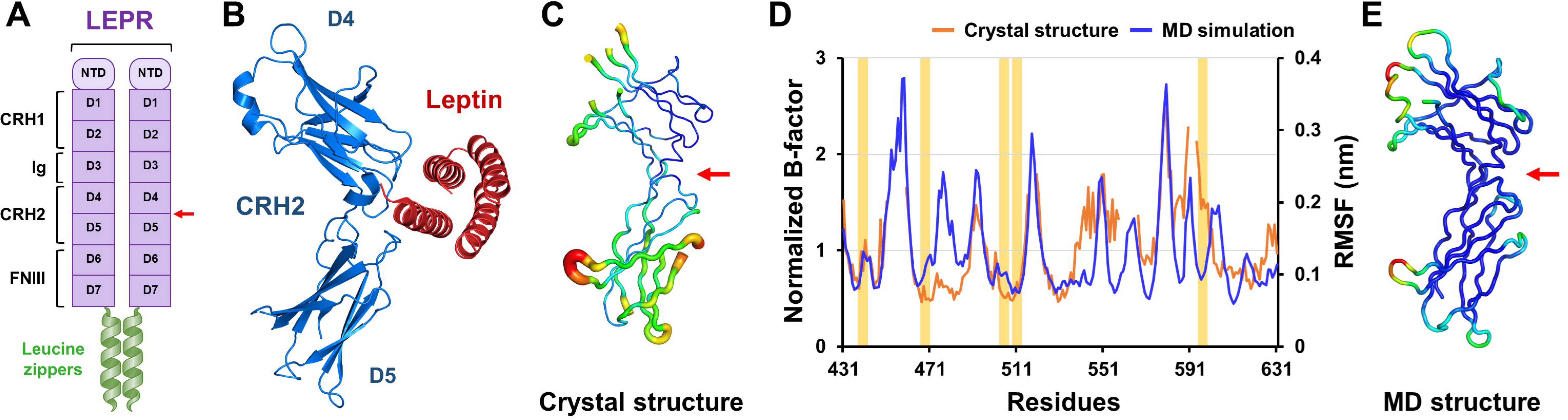
Structural analysis and MD simulation of the LEPR CRH2 domain. (A) Schematic representation of the LEPR ECD domain architecture. A red arrow indicates the leptin- binding site. (B) Structural representation of the leptin–LEPR CRH2 complex (PDB code 7Z3Q). (C) B-factor putty representation of the LEPR CRH2 domain (PDB code 3V6O, Chain A). (D) Normalized B-factors of the LEPR CRH2 domain (PDB code: 3V6O, Chain A) and the RMSF profile of the LEPR CRH2 domain calculated from MD trajectories at 300 K. Red arrows indicate the leptin-binding site identified from the leptin–LEPR CRH2 complex structure. (E) RMSF-derived B-factor putty representation of the LEPR CRH2 domain based on MD trajectories.

The crystal structure of the LEPR CRH2 domain in complex with leptin (PDB code 7Z3Q) revealed partially negative surfaces at the interaction interfaces of both the LEPR CRH2 domain and leptin (Supplementary Figure S1). The interface between the LEPR CRH2 domain and leptin was stabilized by hydrogen bond interactions (Table S1).

To examine the intrinsic dynamics of the leptin-binding site within the LEPR CRH2 domain, we analyzed the B-factors of the LEPR CRH2 domain in its apo (PDB code: 3V6O) and leptin-bound (PDB code: 7Z3Q) states (Figures 1C and Supplementary Figure S2). In the apo state of LEPR CRH2 domain structure, the leptin-binding site exhibited a relatively rigid conformation, whereas loops near the binding pocket (chain A: Lys560–Glu565 and Val589– Lys594) lacked visible electron density and were therefore not modeled, suggesting high flexibility (Figure 1C). In contrast, in the leptin-bound structure, not only the leptin-binding site but also the surrounding loops exhibited low B-factors, suggesting that leptin binding to the LEPR CRH2 domain stabilizes the local structural environment of the leptin-binding region (Supplementary Figure S2). These structural analyses indicate that the central leptin- binding pocket remains rigid and preorganized even in the absence of leptin.

To determine whether this preorganized rigidity was not an artifact of the cryogenic temperatures used for crystallography [51], we performed all-atom MD simulations of the CRH2 domain. RMSF analysis indicated that the leptin-binding site in LEPR remained relatively rigid (Figure 1D and 1E), indicating that the core leptin-binding interface is intrinsically stable. Consequently, we selected this rigid, preorganized cleft between the R4 and R5 subdomains of the LEPR CRH2 domain as the target site for the computational design of miniprotein binders aimed at achieving stable, high-affinity receptor binding.

### 3.2. Computational design of LEPR-binding miniproteins

To design *de novo* miniproteins targeting the LEPR CRH2 domain, an integrated workflow combining hierarchical computational screening with stepwise experimental validation was established (Figure 2A). A parallel *in vitro* screening cascade was implemented to validate the computationally designed miniproteins (Figure 2B). The five selected miniproteins (Mini- 9022, Mini-10457, Mini-12198, Mini-12264, and Mini-19484) were expressed in *E. coli* and purified. Their secondary structures, thermal stability, and protease resistance were then evaluated. The binding affinities of these miniproteins for the LEPR CRH2 domain were measured using MST. Finally, the predicted structures of the LEPR–miniprotein complexes were analyzed, and the predicted binding modes were further evaluated using site-directed mutagenesis and MST.

**Figure 2.**
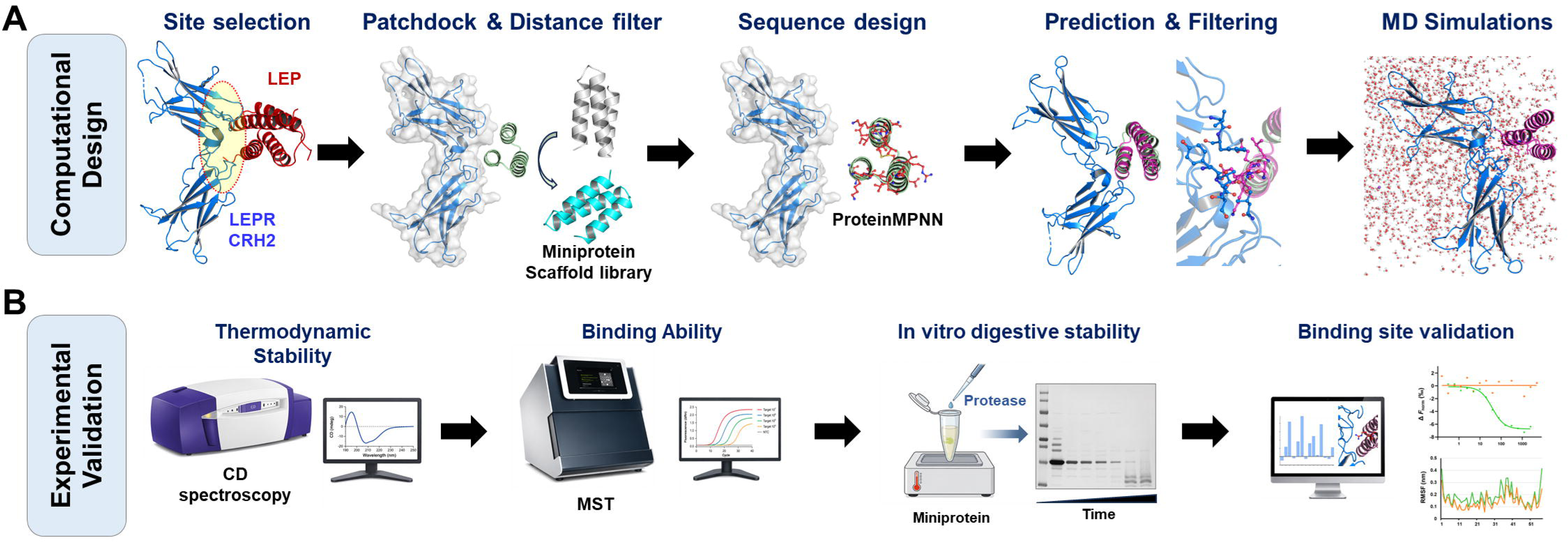
Design and screening workflow for *de novo* miniprotein design and experimental validation. (A) Hierarchical *in silico* design workflow, including target-site selection, PatchDock-based shape-complementarity screening of scaffolds, ProteinMPNN-based sequence redesign, structure prediction using AlphaFold2, and structural filtering using MD simulations. (B) Experimental validation workflow, including characterization of secondary structure, thermal stability, and proteolytic stability; evaluation of binding affinity for the LEPR CRH2 domain using MST; and validation of the predicted binding model through site- directed mutagenesis.

Based on the structural analysis of the LEPR CRH2 domain, we targeted a relatively rigid region within the leptin-binding site for miniprotein design. Because flexible loops adjacent to the leptin-binding site can affect miniprotein scaffold accessibility and influence initial scaffold selection, we used the crystal structure of the LEPR CRH2 domain (chain A, PDB ID 3V6O) as the template. In this structure, the flexible loops adjacent to the leptin-binding site are not modeled.

For *de novo* miniprotein design, the pipeline began with large-scale shape- complementarity screening using PatchDock, in which a library of 20,000 *de novo* miniprotein scaffolds was systematically docked against the target interface of the LEPR CRH2 domain. Based on PatchDock scores and distance-to-target parameters, the initial scaffold pool was narrowed to the top 100 complexes for subsequent sequence optimization. The backbone coordinates of these 100 candidate complexes were then used as fixed templates for sequence redesign with ProteinMPNN, which optimized interfacial residues while preserving the overall scaffold structures.

To evaluate the structural fidelity and foldability of the redesigned sequences, their three- dimensional structures were predicted using AlphaFold2 and aligned with the corresponding design templates (Supplementary Figure S3). The resulting backbone RMSD values ranged from 13.5 to 15.8 Å Given the similarly low RMSD values across the predicted models, the 30 selected miniprotein–LEPR complexes were subjected to 10-ns MD simulations at 300 K as an initial assessment of dynamic stability. During these simulations, the RMSD values of the candidate complexes ranged from 0.31 to 0.49 nm. Based on the 10-ns MD simulations and predicted binding affinities, the five top-ranked miniprotein–LEPR complexes (Mini-9022, Mini-10457, Mini-12198, Mini-12264, and Mini-19484) were selected. Mini-9022, Mini-10457, Mini-12198, Mini-12264, and Mini-19484 consisted of 52, 55, 58, 52, and 57 amino acids, respectively, and showed no sequence similarity among them (Table S2). These miniproteins exhibited typical α-helical bundle structures consisting of three α-helices, similar to those of other miniproteins, whereas their binding positions on the LEPR CRH2 domain differed (Supplementary Figure S4). Given that the initial 10-ns simulations provided only a preliminary assessment of complex stability, the five selected CRH2-targeting miniprotein candidates were subjected to extended 200-ns MD simulations to evaluate longer-term dynamic stability. RMSD analysis showed that Mini-9022, Mini-10457, Mini-12198, and Mini- 19484 maintained relatively stable RMSD profiles within 0.20–0.60 nm throughout the simulations (Supplementary Figure S5). In contrast, the Mini-12264 complex exhibited a sudden, marked increase in RMSD at approximately 115 ns, followed by a higher plateau fluctuating between 0.60 and 0.90 nm. Analysis of the minimum distance between LEPR and each miniprotein throughout the simulations showed that the stable complexes (Mini-9022, Mini-10457, Mini-12198, and Mini-19484) maintained relatively constant distances of 0.14– 0.24 nm, whereas the distance for Mini-12264 increased moderately from approximately 0.14 to 0.28 nm. Although this relatively small change indicates that Mini-12264 did not fully dissociate into the solvent, it suggests a substantial weakening of close interfacial contacts. To further characterize this dynamic instability, representative Mini-12264 structures extracted from two distinct equilibrium phases at 80 and 180 ns were superimposed (Supplementary Figure S5). The structural alignment showed substantial spatial displacement and interfacial sliding of Mini-12264 relative to the rigid, preorganized LEPR binding pocket (Supplementary Figure S5).

### 3.3. Validation of miniprotein binding affinity and thermostability

To determine whether the *de novo*-designed miniproteins (Mini-9022, Mini-12264, Mini- 12198, Mini-10457, and Mini-19484) were properly folded and able to bind LEPR, we examined their secondary structures, thermal stability, and binding affinity for the LEPR CRH2 domain.

CD spectroscopy showed that Mini-9022, Mini-10457, Mini-12198, and Mini-19484 exhibited characteristic α-helical spectra at 25 °C, with distinct minima at 208 and 222 nm (Figure 3A and Table S3). In contrast, Mini-12264 exhibited a minimum near 200 nm, consistent with a predominantly disordered or random-coil conformation (Figure 3A), indicating that Mini-12264 did not adopt the intended folded state despite the stable structure predicted by AI-based modeling and MD simulations.

**Figure 3.**
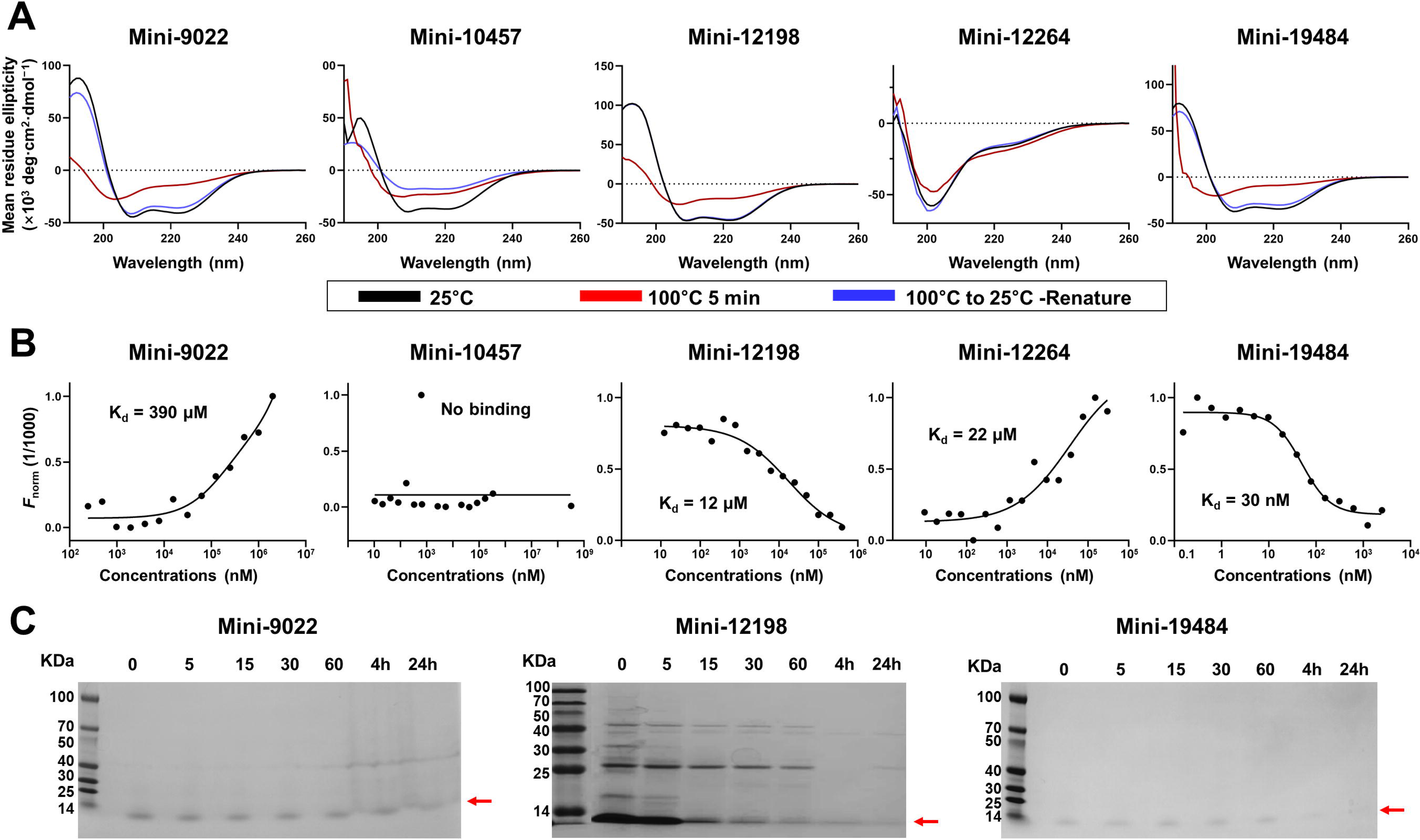
Validation of recombinant *de novo*-designed miniproteins. (A) Circular dichroism spectroscopy analysis of the secondary structure and thermal stability of Mini-9022, Mini- 10457, Mini-12198, Mini-12264, and Mini-19484 under three conditions: initial state at 25 °C (black), after 5 min of thermal denaturation at 100 °C (red), and following refolding upon cooling to 25 °C (blue). (B) MST analysis of Mini-9022, Mini-10457, Mini-12198, Mini-12264, and Mini-19484 binding to the LEPR CRH2 domain. (C) Time-dependent protease digestion assay of Mini-9022, Mini-12198, and Mini-19484 using pepsin.

Thermal refolding analysis is important for evaluating whether *de novo*-designed miniproteins can reproducibly recover their intended conformations after thermal denaturation [29, 30]. To further assess the folding robustness of the designed miniproteins, thermal denaturation at 100 °C for 5 min followed by cooling to 25 °C was analyzed using CD spectroscopy. After thermal denaturation at 100 °C, Mini-10457, Mini-12264, and Mini- 19484 exhibited spectra characteristic of predominantly disordered conformations, whereas Mini-9022 and Mini-12198 retained partial α-helical content accompanied by increased random-coil characteristics (Figure 3A). Following cooling to 25 °C, Mini-12198 recovered a

CD spectrum similar to that observed before heating, indicating efficient and reversible folding into the designed α-helical bundle (Figure 3A). Mini-9022 and Mini-19484 exhibited weaker α-helical signals after refolding than in their original spectra, suggesting partial loss of the native α-helical bundle structure during the thermal cycle (Figure 3A). Mini-10457 showed a further reduction in α-helical content after refolding, indicating poor structural stability during thermal denaturation and recovery (Figure 3A). Meanwhile, Mini-12264 remained predominantly disordered throughout the experiment, suggesting that this miniprotein does not favor stable α-helical folding even after thermal refolding.

Next, to characterize interactions between the miniproteins and the LEPR CRH2 domain, binding affinities were measured using MST. Mini-19484 exhibited a dose-dependent binding curve with a *K*_d_ of 30 nM (Figure 3B), indicating that it is a promising lead binder for LEPR CRH2. Mini-12198 and Mini-12264 showed moderate binding affinities (*K*_d_ = 12 μM and 22 μM), respectively, whereas Mini-9022 showed substantially weaker binding (*K*_d_ = 390 μM), suggesting weak or potentially nonspecific interactions. In contrast, Mini-10457 exhibited no detectable binding despite the stable binding behavior predicted by the MD simulations.

Notably, the LEPR CRH2-binding miniproteins exhibited two distinct ligand-dependent MST response patterns (Figure 3B). The negative MST responses observed for Mini-12198 and Mini-19484 may reflect altered solvation properties or local conformational rearrangements induced by complex formation. In contrast, the positive MST responses observed for Mini-9022 and Mini-12264 may indicate reduced conformational flexibility and decreased hydration upon ligand binding. Similar bidirectional MST responses have been reported and attributed to differences in thermophoretic mobility, hydration-shell properties, charge distribution, and fluorophore environment rather than to experimental artifacts [52].

Protease resistance is an important characteristic for evaluating the structural stability of designed miniproteins, as enhanced resistance to proteolytic degradation is closely associated with compact folding, improved durability, and potential biotechnological or therapeutic applicability [53]. The proteolytic stability of Mini-9022, Mini-12198, and Mini- 19484, which showed potential as LEPR binders, was further examined using pepsin digestion assays. The other miniproteins were excluded because of unintended protein folding and a lack of binding affinity for LEPR. In the pepsin stability assay, Mini-12198 was rapidly digested, with the majority of the protein degraded by 60 min. In contrast, Mini-9022 and Mini-19484 exhibited superior resistance to proteolytic cleavage, remaining largely intact at 60 min. Although both Mini-9022 and Mini-19484 exhibited partial degradation at 4 h and Mini-19484 was completely degraded by 24 h, this eventual clearance provides a pharmacological advantage by minimizing the risks of long-term in vivo accumulation and immunotoxicity. (Figure 3C). Taken together, these findings indicate that Mini-19484 exhibited stable protein folding, high binding affinity for LEPR CRH2, and remarkable resistance to proteolytic degradation, suggesting that it is the most promising LEPR CRH2- binding miniprotein.

### 3.4. Identification of Key Binding Hotspots using Mutational Analysis

The high binding affinity of Mini-19484 for the LEPR CRH2 domain suggests that it may serve as a promising scaffold for developing future therapeutics or molecular probes targeting LEPR. To gain structural insights into miniprotein engineering aimed at modulating binding affinity, the molecular interactions between Mini-19484 and the LEPR CRH2 domain were analyzed.

In the structural model of the Mini-19484–LEPR complex, the binding interface was stabilized by 10 hydrogen bonds and 6 salt bridges (Figure 4A and Table S4). Among these interactions, the NH1, NH2, and NE atoms of Arg18 in Mini-19484 formed three hydrogen bonds and six salt bridges with the OH atom of Tyr617 and the OD1 and OD2 atoms of Asp532 and Asp617 in CHR2 domain of LEPR. To quantitatively evaluate the energetic contributions of these residues, computational alanine-scanning mutagenesis was performed. Among the analyzed residues, the R18A mutation produced the highest predicted destabilization energy (ΔΔG = 2.4 kcal/mol) (Figure 4B), suggesting that Arg18 functions as a critical binding hotspot. To further elucidate the structural impact of this critical residue, MD simulations were performed on the Mini-19484-R18A mutant in complex with LEPR. RMSF analysis revealed a marked decrease in the flexibility across residues 11–20 following the R18A mutation. Notably, the RMSF value at position 18 decreased by more than 0.1 nm, indicating significant local stabilization.(Supplementary Figure S6). The number of interfacial hydrogen bonds formed in the Mini-19484–LEPR and Mini-19484-R18A–LEPR complexes was monitored over the 200-ns simulation trajectories (Supplementary Figure S7). Compared with the Mini-19484–LEPR complex, the Mini-19484-R18A mutant exhibited fewer hydrogen bonds, suggesting that Arg18 may greatly influence LEPR binding. To validate the computational analysis, an *in vitro* binding analysis was subsequently performed using MST. The Mini-19484-R18A mutant completely lost detectable binding to LEPR (Figure 4C), indicating that the Arg18 side chain is critical for the Mini-19484–LEPR interaction.

**Figure 4.**
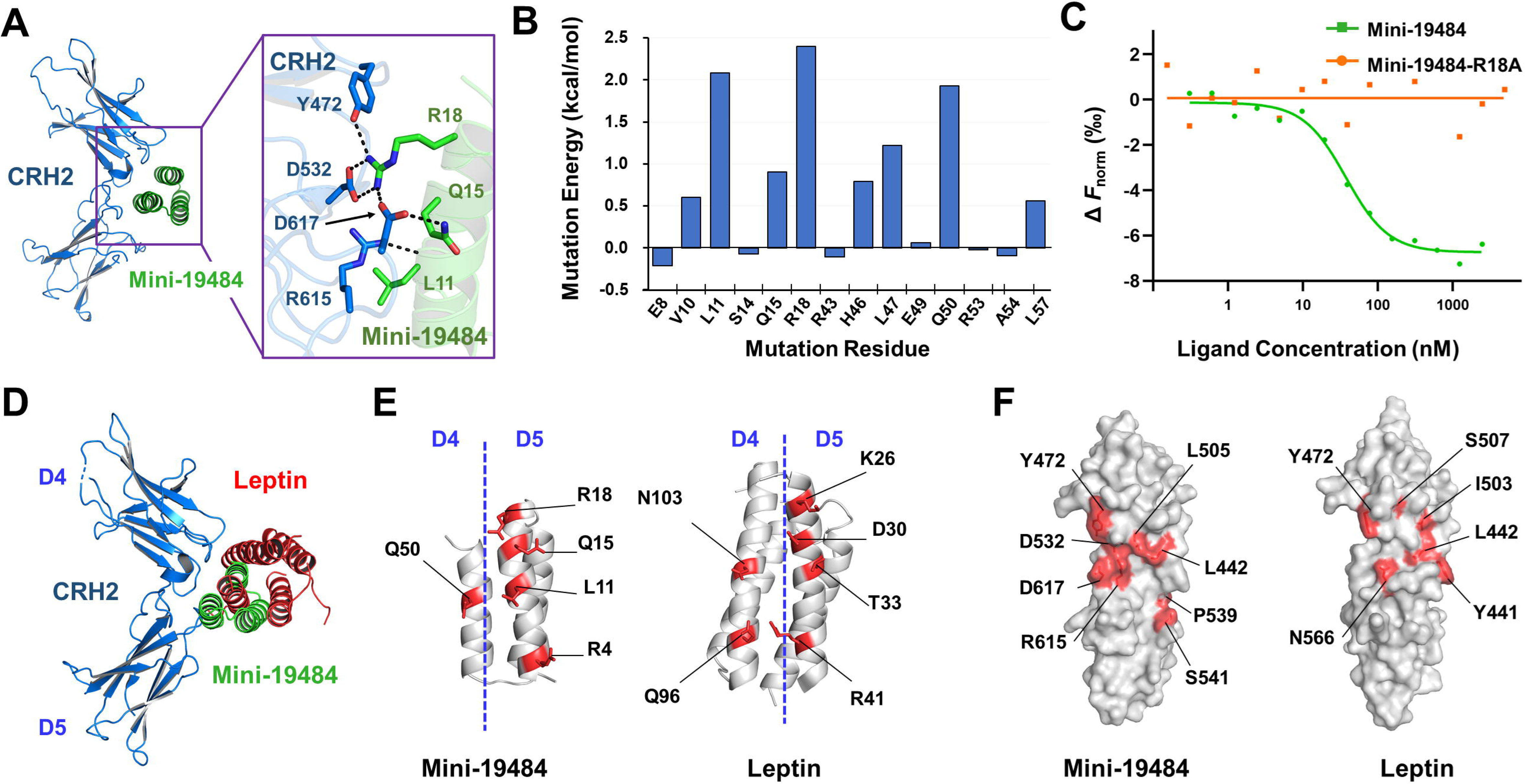
Identification and validation of key molecular hotspots involved in LEPR recognition. (A) Structural model of Mini-19484 bound to LEPR CRH2 and corresponding interaction analysis of the binding interface. (B) Computational alanine-scanning mutagenesis of Mini-19484 residues involved in LEPR-CRH2 binding, identifying Arg18 as the critical energetic hotspot residue. (C) MST analysis of Mini-19484 (green) and the Mini- 19484-R18A mutant (orange) binding to the LEPR CRH2 domain. (D) Superimposition of LEPR CRH2-bound Mini-19484 (green) and leptin (red) structures. (E) Interaction interfaces of Mini-19484 and leptin (PDB code 7Z3Q, chain E) with the LEPR CRH2 domain. (F) Interaction interfaces of the LEPR CRH2 domain with Mini-19484 and leptin (PDB code 7Z3Q, chain F). Interacting residues are highlighted in red and shown as surface and stick representations.

To test whether Mini-19484 exhibits binding characteristics comparable to those of leptin toward LEPR CRH2, the predicted Mini-19484–LEPR complex was structurally compared with the leptin–LEPR structure (PDB code 7Z3Q) (Figure 4D). Mini-19484 consists of three α-helices arranged in a triangular architecture, with the α2 helix inserted into the cleft between the two CRH2 domains. In contrast, leptin, which comprises four α-helices, primarily interacts with the surface of the CRH2 domains without penetrating deeply into the cleft between them. Structural comparison showed that Mini-19484 was positioned more deeply within the cleft between the two CRH2 domains than leptin, indicating a distinct binding mode (Figure 4D). The buried surface area of Mini-19484 and leptin was approximately 785 and 830 Å^2^, respectively. The calculated binding energies of Mini-19484 and leptin were approximately −11.3 and −12.0 kcal/mol, respectively, indicating comparable predicted binding energies, with a slightly more favorable value for leptin. This indicates that, despite its considerably smaller size, Mini-19484 exhibits a buried surface area comparable to that of leptin and forms an extensive interaction interface with the LEPR CRH2 domain. Structural analysis showed that both Mini-19484 and leptin interact with the D4 and D5 domains of LEPR CRH2, but their binding positions differ. The binding position of Mini-19484 is shifted toward the D5 domain, whereas leptin is positioned more centrally between the D4 and D5 domains (Figure 4E).

Furthermore, the LEPR surface residues involved in binding to Mini-19484 in the predicted model and those involved in leptin binding in the crystal structures exhibited distinct spatial distributions, although two interacting residues (Leu442 and Tyr472) were shared. In the Mini-19484–LEPR model, the interacting residues (Leu442, Tyr472, Leu505, Asp532, Pro539, Ser541, Arg615, and Asp617) were distributed across two spatially distinct regions. In contrast, the leptin-interacting residues (Tyr441, Leu442, Tyr472, Ile503, Ser507, and Asn566) in the crystal structures were more broadly distributed across the central region of the LEPR surface (Figure 4F). Collectively, these results indicate that Mini-19484 recognizes LEPR CRH2 through a binding mode distinct from that of leptin, characterized by deeper insertion into the CRH2 cleft while maintaining an extensive interaction interface.

## 4. Discussion

LEPR plays a central role in regulating appetite and energy homeostasis [16, 54]. Modulating LEPR signaling or leptin–LEPR interactions may provide a potential therapeutic strategy for various metabolic disorders, including obesity.

In this study, we showed that *de novo* computational protein design can generate miniprotein binders targeting the leptin-binding site within the LEPR CRH2 domain. Through hierarchical computational screening and biochemical characterization, three miniproteins— Mini-9022, Mini-12198, and Mini-19484—were identified as binders of LEPR CRH2.

Recent advances in computational protein design, particularly artificial intelligence–based approaches for structure prediction and sequence design, have greatly expanded the ability to generate miniprotein binders with predefined structural and binding properties [33, 55]. Nevertheless, the computational predictions in this study were partially consistent with the experimental results. For example, computational modeling predicted Mini-12264 to adopt a stable α-helical structure, whereas the recombinant protein predominantly exhibited disordered or random-coil characteristics in CD spectroscopy. In addition, Mini-10457 maintained a stable interaction with LEPR CRH2 during the MD simulations but showed no detectable binding in the MST assay. These discrepancies highlight a limitation of current computational protein design approaches, as predicted structural stability and binding properties do not always translate into correct folding or target binding under experimental conditions. Therefore, experimental validation remains essential for evaluating computationally designed miniproteins.

Our biochemical characterization showed that Mini-9022, Mini-12198, and Mini-19484 formed stable α-helical structures and bound to the LEPR CRH2 domain. Among these miniproteins, Mini-19484 showed nanomolar binding affinity for LEPR CRH2 (*K*_d_ = 30 nM) and showed no detectable degradation after 24 h of pepsin treatment under the tested conditions. These properties are consistent with those reported for successfully designed *de novo* miniproteins, which exhibit high binding affinity and structural stability [30]. Although Mini-9022 and Mini-12198 showed relatively weaker binding affinities and lower proteolytic stability than Mini-19484, they may also serve as scaffolds for further engineering to improve LEPR binding and proteolytic stability.

Computational alanine scanning, site-directed mutagenesis, and MST analysis identified Arg18 of Mini-19484 at the predicted interface as a key residue contributing to LEPR binding. However, these mutagenesis results support the importance of Arg18 in LEPR binding but do not confirm the precise binding interface of the predicted LEPR–Mini-19484 complex. Therefore, experimental determination of the LEPR–Mini-19484 complex structure is necessary to define the molecular interactions between LEPR and Mini-19484. Such structural information could provide a basis for further engineering of Mini-19484 to improve its binding affinity for LEPR.

Notably, structural analysis revealed that the predicted binding mode of Mini-19484 differed substantially from that of natural leptin. Natural leptin interacts primarily with the surface of the LEPR CRH2 domain, whereas Mini-19484 was predicted to bind more deeply within the CRH2 domain cleft. Thus, the predicted binding architecture of Mini-19484 does not directly mimic the molecular interaction mode of natural leptin. Although Mini-19484 exhibited nanomolar binding affinity for the LEPR CRH2 domain (*K*_d_ = 30 nM), its affinity was weaker than that reported for natural leptin (*K*_d_ = 0.2–15 nM, depending on the experimental method) [56, 57]. Therefore, although Mini-19484 binds the leptin-binding region of LEPR CRH2, its lower affinity suggests that further optimization may be needed for effective competition with native leptin. Nevertheless, Mini-19484 may provide a useful structural framework for the further development of LEPR-targeting miniproteins with improved binding affinity. Future studies should determine whether engineered Mini-19484 variants can effectively compete with native leptin for LEPR binding and evaluate their effects on LEPR- mediated cellular signaling.

## 5. Conclusion

In this study, we report *de novo* miniproteins that recognize the leptin-binding region of LEPR using an integrated computational and experimental approach. The discrepancies between computational predictions and experimental characterization emphasize the importance of biochemical validation in *de novo* protein design. Among the designed miniproteins, Mini-19484 exhibited the most favorable combination of structural stability, proteolytic resistance, and LEPR-binding affinity, with a *K*_d_ of 30 nM. These findings establish Mini-19484 as a useful starting scaffold for further engineering of LEPR-targeting miniproteins with improved binding properties.

## Acknowledgments

The National Supercomputing Center supported this work by providing supercomputing resources and technical support. During manuscript preparation, the authors used ChatGPT 5.2 for minor English-language corrections and adjustments. After using this tool, the authors reviewed and edited the content as needed and took full responsibility for the publication.

## CRediT authorship contribution statement

**Xue Bai**: Investigation, Methodology, Data curation, Visualization, Formal analysis, Writing – original draft. **Zhidan Zhao**: Methodology, Data curation, Visualization, Formal analysis. **Yu Jin**: Methodology, Data curation. **Jianyun Guo**: Data curation. **Nam-Chul Ha**: Data curation, Visualization, Formal analysis. **Ki Hyun Nam**: Formal analysis, Funding acquisition, Data curation, Visualization, Writing – original draft, Writing – review & editing. **Yongbin Xu**: Conceptualization, Investigation, Methodology, Resources, Funding acquisition, Writing – original draft, Writing – review & editing.

## Data availability

The data that support the findings of this study are available from the corresponding author upon reasonable request.

