## Supplementary Data for "*De Novo* Design of Miniprotein Binders Targeting the Leptin-Binding Site of the Leptin Receptor"

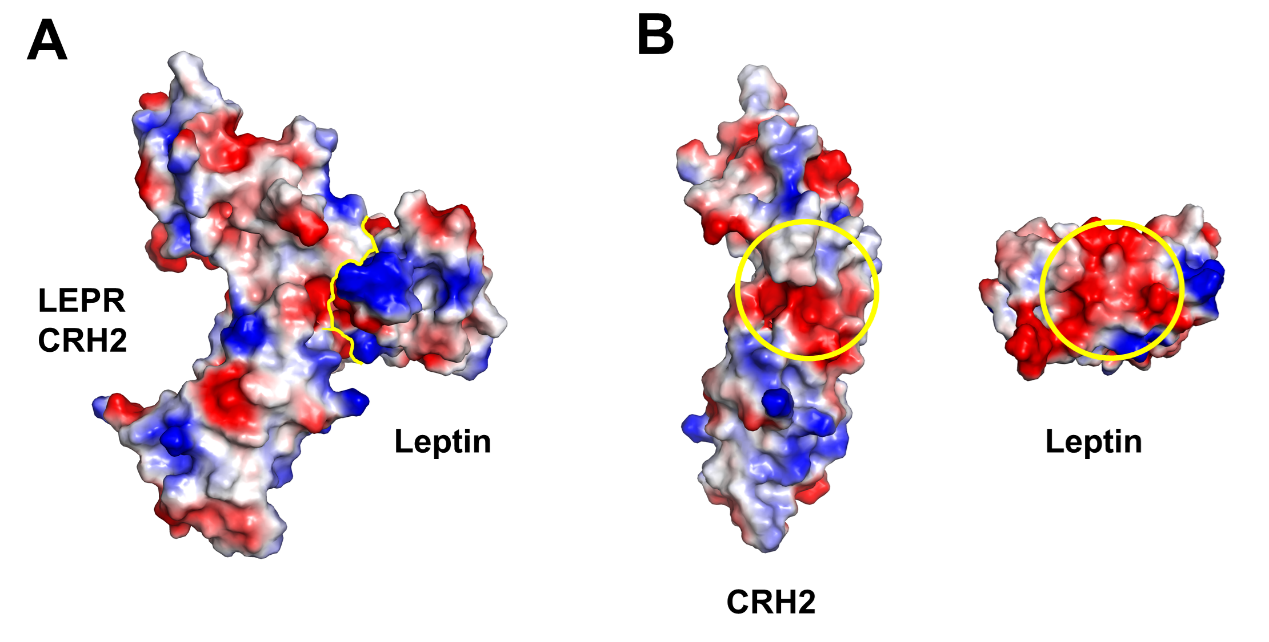

**Figure S1**. (A) Electrostatic potential surface representation of the LEPR CRH2 domain in complex with leptin (PDB code: 7Z3Q). Yellow indicates the interaction interface between the LEPR CRH2 domain and leptin. (B) Interaction interface between the LEPR CRH2 domain and leptin. The buried surface area of the LEPR CRH2 domain upon leptin binding is highlighted by yellow circles.

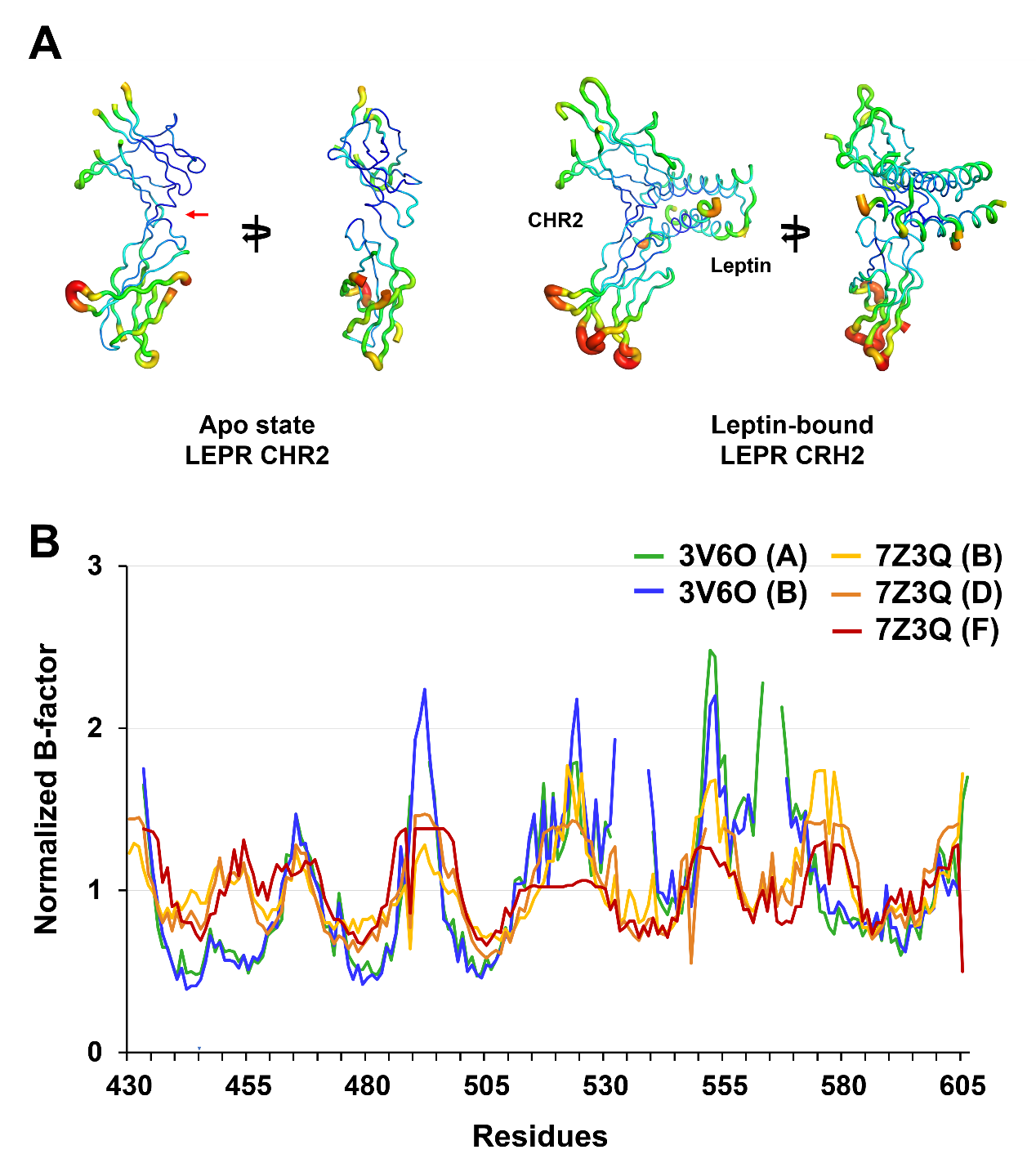

**Figure S2.** B-factor analysis of the LEPR CRH2 domain in the apo (PDB code: 3V6O) and leptin-bound (PDB code: 7Z3Q) states. (A) B-factor putty representations and normalized B-factor profiles of the LEPR CRH2 domain in the apo and leptin-bound states. (B) Normalized B-factor profiles of LEPR in the apo (PDB code: 3V6O) and leptin-bound (PDB code: 7Z3Q) states.

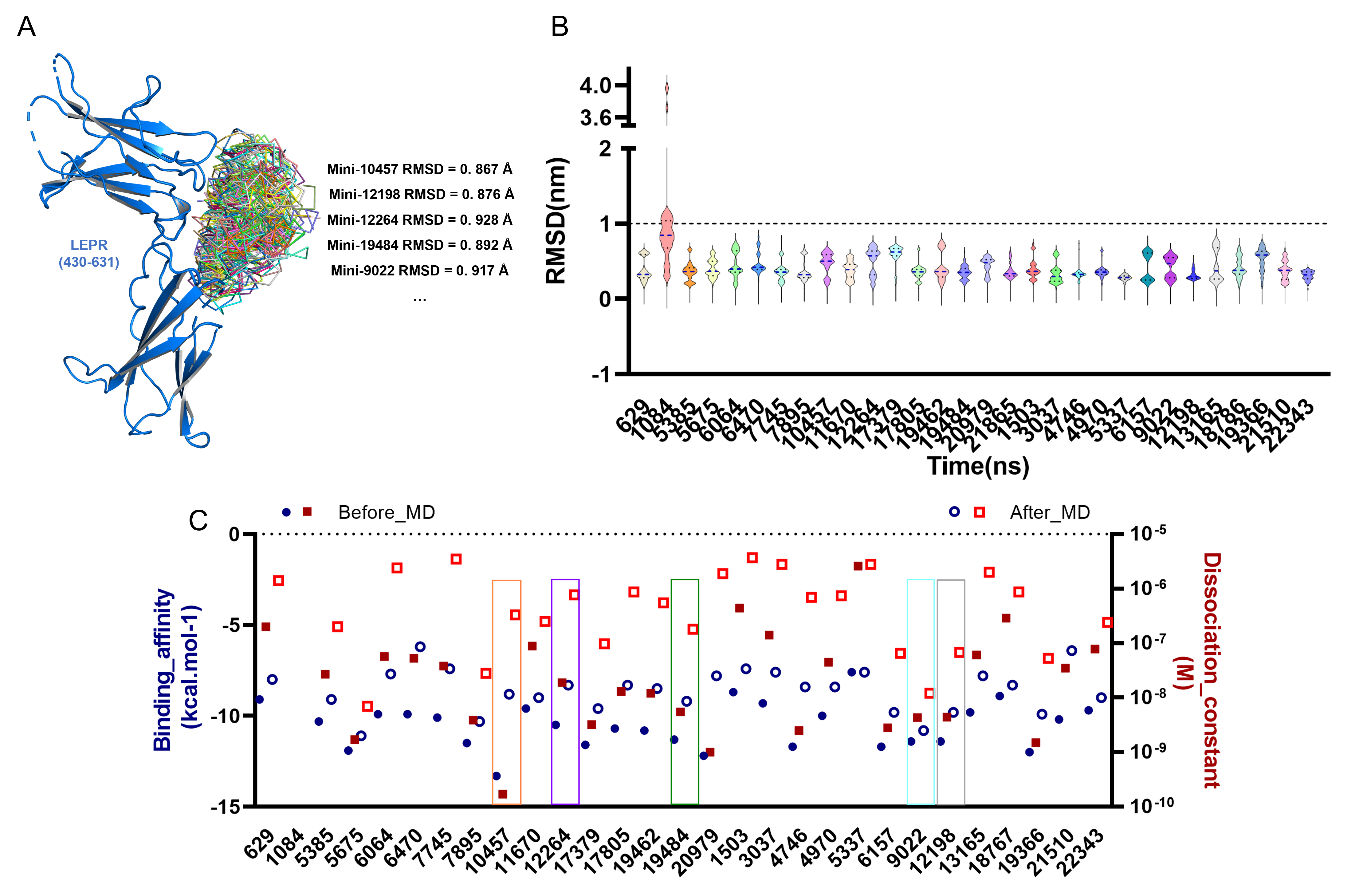

**Figure S3.** *In silico* folding validation and initial 10-ns molecular dynamics screening of redesigned scaffolds. (A) Structural superposition of AlphaFold2-predicted models with their respective PatchDock templates, confirming preservation of the predicted binding modes (RMSD < 1.0 Å). (B) Violin plots showing the RMSD distributions during the 10-ns molecular dynamics (MD) simulations. (C) Binding affinities predicted by PRODIGY after the MD simulations and used to prioritize the top candidates.

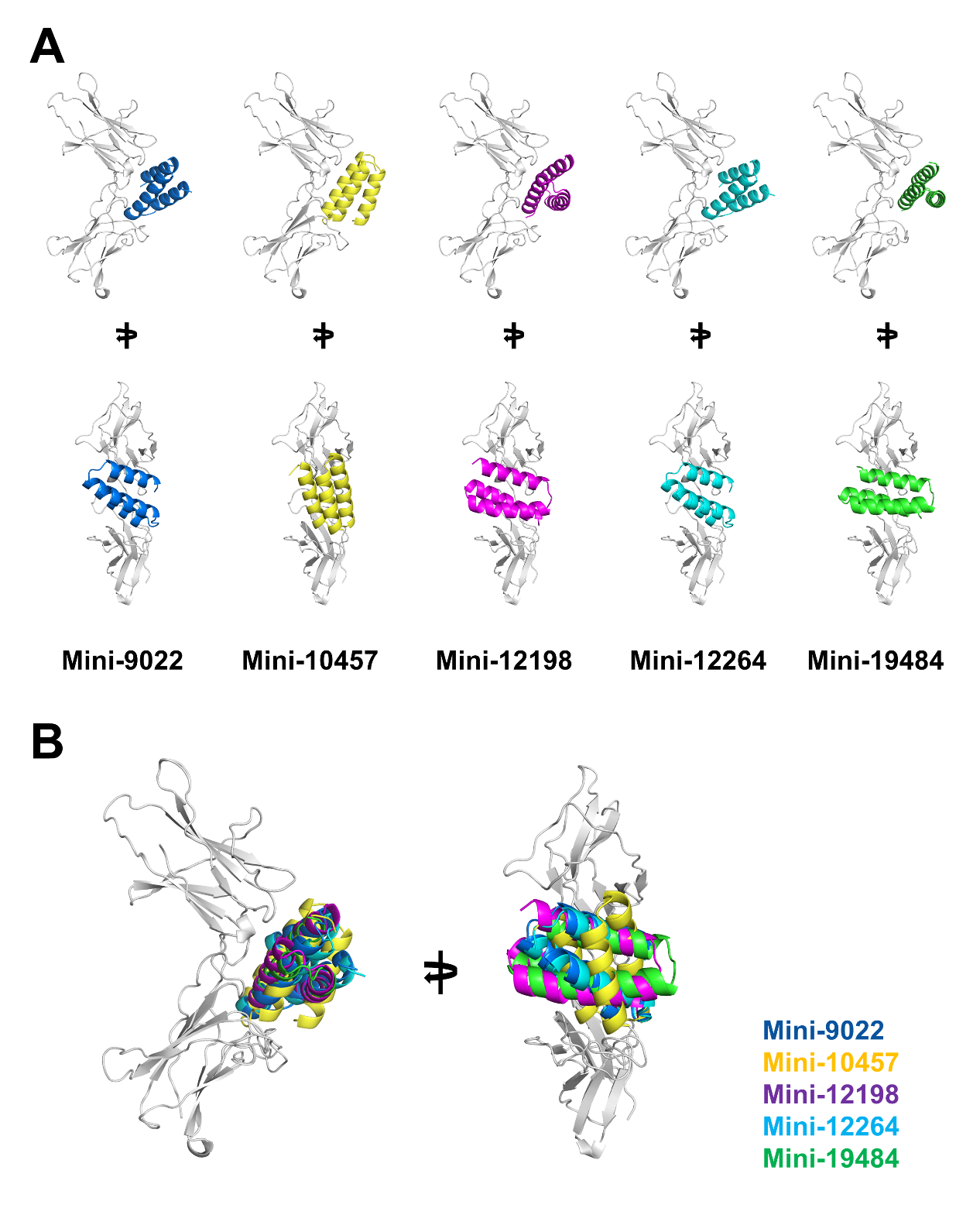

**Figure S4**. Cartoon representations of (A) the selected miniproteins (Mini-9022, Mini-10457, Mini-12198, Mini-12264, and Mini-19484) and (B) their predicted binding models with the LEPR CRH2 domain.

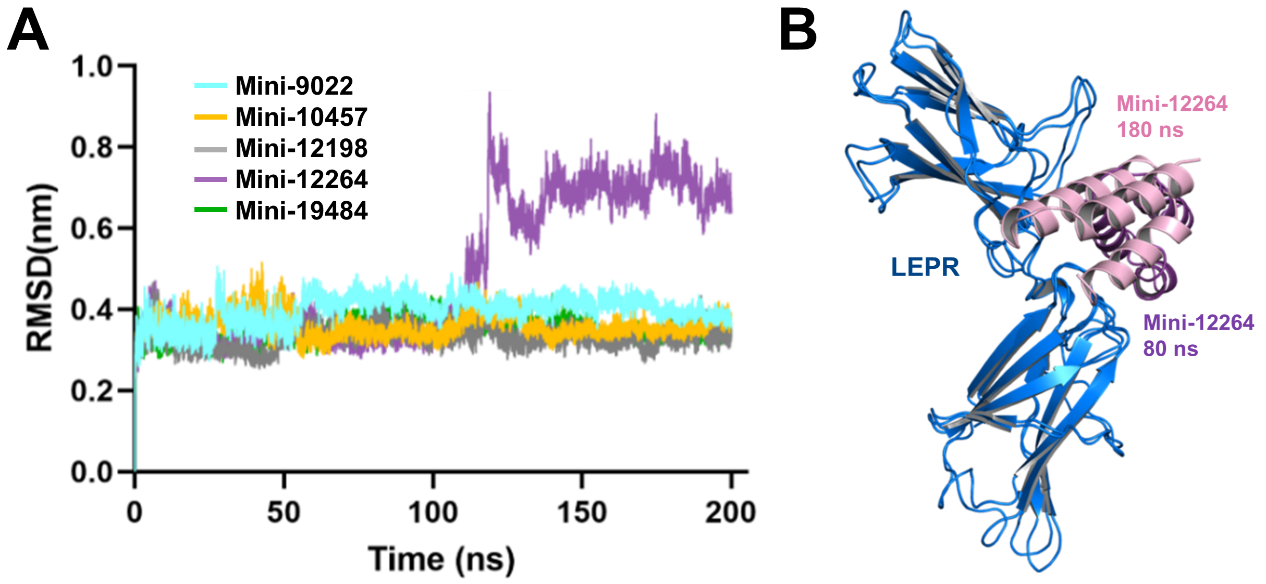

**Figure S5.** MD simulations of LEPR complexes with the designed miniproteins Mini-9022, Mini-10457, Mini-12198, Mini-12264, and Mini-19484. (A) RMSD profiles of the miniprotein–LEPR complexes during 200-ns MD simulations. (B) Structural superimposition of the LEPR–Mini-12264 complex at different time points during the MD simulation.

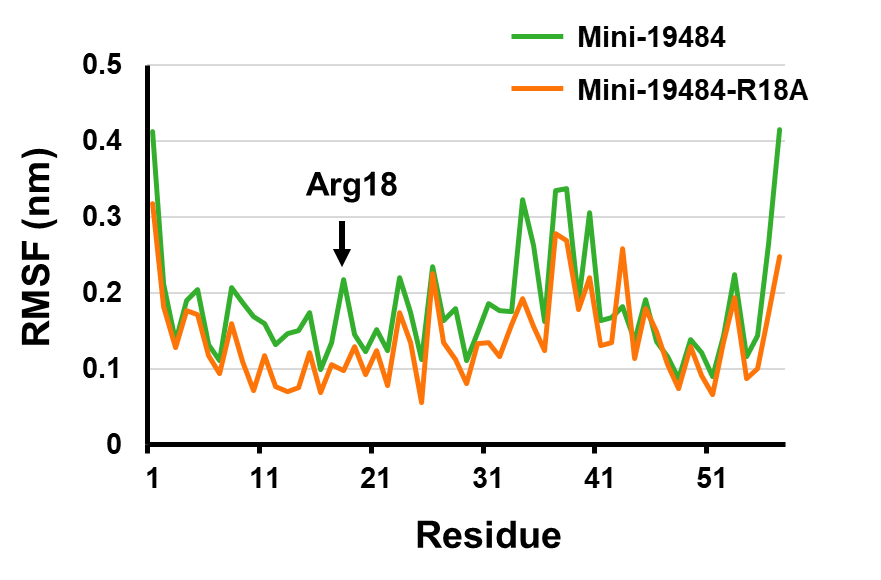
**Figure S6**. RMSF profiles of the LEPR–Mini-19484 (green) and LEPR–Mini-19484-R18A (yellow) complexes obtained from MD simulations.

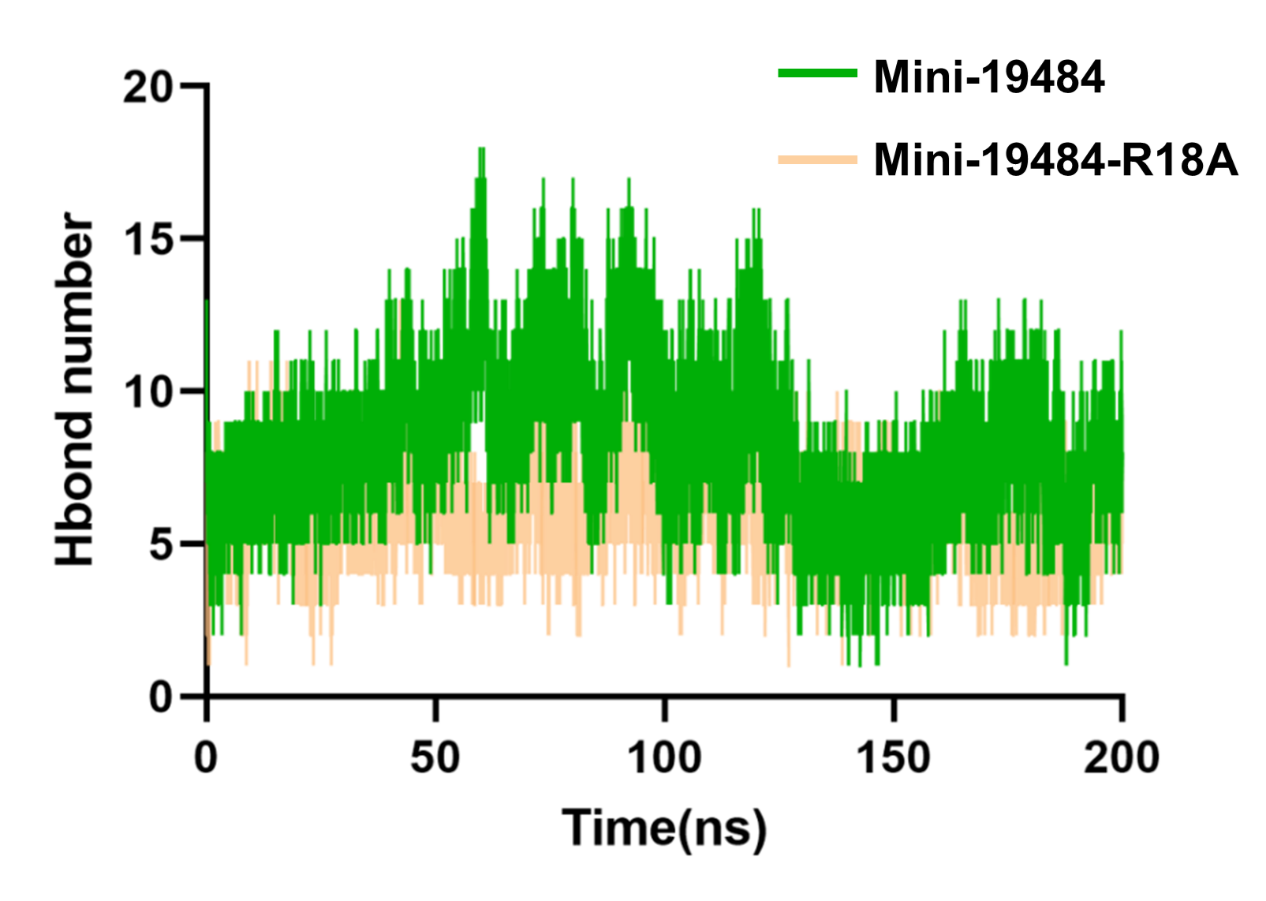

**Figure S7.** Number of hydrogen-bond interactions between Mini-19484 (green) or Mini-19484-R18A (yellow) and LEPR during the 200-ns MD simulation

**Table S1**. Hydrogen bond interactions between the LEPR CRH2 domain and leptin in the crystal structure (PDB code: 7Z3Q).

| **LEPR CHR2 (Chain B)** | **Distance (Å)** | **Leptin (chain C)** |
| --- | --- | --- |
| Tyr441 [O] | 2.97 | Arg41 [NH2] |
| Leu442 [O] | 2.83 | Arg41 [NH1] |
| Tyr472 [OH] | 2.77 | Asp30 [OD2] |
| Ile503 [O] | 3.84 | Asn99 [ND2] |
| Ile503 [O] | 2.88 | Gln96 [NE2] |
| Ile503 [N] | 3.22 | Gln96 [OE1] |
| **LEPR CHR2 (Chain D)** | **Distance (Å)** | **Leptin (chain A)** |
| Leu442 [O] | 2.45 | Arg41 [NH1] |
| Tyr472 [OH] | 3.10 | Asp30 [OD2] |
| Ile503 [N] | 3.22 | Gln96 [OE1] |
| Ile503 [O] | 2.88 | Gln96 [NE2] |
| Ser507 [OG] | 3.36 | Asn103 [OD1] |
| Asn566 [ND2] | 3.60 | Thr33 [OG1] |
| **LEPR CHR2 (Chain F)** | **Distance (Å)** | **Leptin (chain E)** |
| Tyr441 [O] | 3.35 | Arg 41 [NH2] |
| Leu442 [O] | 2.92 | Arg 41 [NH1] |
| Tyr472 [OH] | 3.50 | Lys26 [O] |
| Tyr472 [OH] | 3.52 | Asp30 [OD2] |
| Ile503 [N] | 3.43 | Gln96 [OE1] |
| Ile503 [O] | 3.03 | Gln96 [NE2] |
| Ser507 [OG] | 3.59 | Asn103 [OD1] |
| Asn566 [ND2] | 3.60 | Thr33 [OG1] |

**Table S2.** Amino acid sequences of the designed miniproteins

| Name | Length | Sequence |
| --- | --- | --- |
| Mini-9022 | 52 | NAVEKVKELCEKALKIKDPTERLELIFEALDICRESNSLECARILNETAKKL |
| Mini-10457 | 55 | SSRALEFAKRIIGILKGETSETARTAIYNNSKKQLEALNNPEALELLEKEAEKIF |
| Mini-12198 | 58 | SLVEESTALLLELSSLLEANKPEEFEKTYKKAKEVIEKLNNPDFKRALQQLEKAFKEK |
| Mini-12264 | 52 | NGFEKVKKLCEEAKKEPDPTERFELLSEARDLAVESKNPEAQKYVEKVAQEL |
| Mini-19484 | 57 | SALREVTELVLAVSQLVRQGKPEEAREKLEEIEKIAEKHNDPRLKHLAEQAKRALEL |

**Table S3.** Relative α-helical contents were estimated qualitatively from the circular dichroism spectra

| Temperature | Mini-9022 | Mini-10457 | Mini-12198 | Mini-12264 | Mini-19484 |
| --- | --- | --- | --- | --- | --- |
| 25°C | **+++++** | **+++++** | **+++++** | **-** | **+++++** |
| 100°C | **-** | **++** | **++** | **-** | **-** |
| 100°C -> 25°C | **++++** | **+++** | **+++++** | **-** | **++++** |

**Table S4**. Interactions between the Mini-19484 and LEPR CRH2 domain in the model structure.

| **Hydrogen bonds** | | |
| --- | --- | --- |
| **LEPR CHR2** | **Distance (Å)** | **Mini-19484** |
| Leu442 [O] | 3.62 | Gln50 [NE2] |
| Tyr472 [OH] | 3.17 | Arg18 [NH1] |
| Leu505 [N] | 3.47 | Gln50 [OE1] |
| Asp532 [OD1] | 3.10 | Arg18 [NH2] |
| Asp532 [OD2] | 2.55 | Arg18 [NH1] |
| Pro539 [O] | 3.69 | Arg4 [NH2] |
| Ser541[OG] | 3.59 | Arg4 [NH2] |
| Arg615 [NH2] | 3.11 | Leu11 [O] |
| Asp617 [OD1] | 3.20 | Gln15 [NE2] |
| Asp617 [OD2] | 2.56 | Arg18 [NH2] |
| **Salt Bridges** | | |
| Asp532 [OD1] | 3.52 | Arg18 [NH1] |
| Asp532 [OD1] | 3.10 | Arg18 [NH2] |
| Asp532 [OD2] | 2.55 | Arg18 [NH1] |
| Asp532 [OD2] | 3.62 | Arg18 [NH2] |
| Asp617 [OD2] | 3.42 | Arg18 [NE] |
| Asp617 [OD2] | 2.56 | Arg18 [NH2] |
